# Beyond the Michaelis-Menten equation: Accurate and efficient estimation of enzyme kinetic parameters

**DOI:** 10.1101/211961

**Authors:** Boseung Choi, Grzegorz A. Rempala, Jae Kyoung Kim

**Affiliations:** Korea University Sejong campus, Division of Economics and Statistics, Department of National Statistics, Sejong, 30019, Korea; The Ohio State University, Division of Biostatistics and Mathematical Biosciences Institute, Columbus, OH 43210, USA; Korea Advanced Institute of Science and Technology, Department of Mathematical Sciences, Daejeon, 34141, Korea

## Abstract

Examining enzyme kinetics is critical for understanding cellular systems and for using enzymes in industry. The Michaelis-Menten equation has been widely used for over a century to estimate the enzyme kinetic parameters from reaction progress curves of substrates, which is known as the progress curve assay. However, this canonical approach works in limited conditions, such as when there is a large excess of substrate over enzyme. Even when this condition is satisfied, the identifiability of parameters is not always guaranteed, and often not verifiable in practice. To overcome such limitations of the canonical approach for the progress curve assay, here we propose a Bayesian approach based on an equation derived with the total quasi-steady-state approximation. In contrast to the canonical approach, estimates obtained with this proposed approach exhibit little bias for any combination of enzyme and substrate concentrations. Importantly, unlike the canonical approach, an optimal experiment to identify parameters with certainty can be easily designed without any prior information. Indeed, with this proposed design, the kinetic parameters of diverse enzymes with disparate catalytic efficiencies, such as chymotrypsin, fumarase, and urease, can be accurately and precisely estimated from a minimal amount of timecourse data. A publicly accessible computational package performing the Bayesian inference for such accurate and efficient enzyme kinetics is provided.

## Introduction

Because enzymes can modulate biochemical reaction rates by selectively catalyzing specific substrates^1^, they play fundamental roles in metabolism, signal transduction, and cell regulation, and their malfunction can cause serious diseases^2, 3^. Furthermore, enzymes have been used as extremely specific catalysts in diverse industrial fields such as drug development, biofuel production, and food processing^4^. A canonical approach used to understand enzyme kinetics for a century has been based on the Michaelis-Menten equation (MM equation), which was developed by Michaelis and Menten^5^ and then was more rigorously derived by Briggs and Haldane^6^ using the standard quasi-steady-state approximation (sQSSA)^7^. The equation describes the dependence of enzyme-catalyzed reaction rates on the concentration of substrate by using two parameters, the catalytic constant, *k*_*cat*_ and the Michaelis-Menten constant, *K*_*M*_ (see below for details). The *k*_*cat*_ determines the maximum rate of the reaction at saturating substrate concentrations, *V*_*max*_ = *k_cat_ E*_*T*_, where *E*_*T*_ is total enzyme concentration, and the *K*_*M*_ is the substrate concentration at which the reaction rate is half of *V*_*max*_.

There are two major assays to estimate *k*_*cat*_ and *K*_*M*_ from a measured accumulation of product over time (i.e. progress curve): the initial velocity assay (initial rate analysis) and the reaction progress curve assay (progress curve analysis)^8–12^. For the initial velocity assay, initial rates of the reaction are measured for a range of substrate concentrations. Then, by using a linear transform of these data, such as Lineweaver-Burk plots, the two parameters can be easily estimated without use of any strong computational tools^8, 9^. Recent advances in computational tools have led to an alternative approach: the reaction progress curve assay. In this assay, the entire timecourse (i.e. progress curve) is fitted to the solution of a differential equation or integrated rate equation, and thus the data is used more efficiently than in the initial velocity assay^10, 11, 13^. Albeit more technically challenging, the progress curve assay requires less data to estimate parameters than the initial velocity assay does.

Since both assays are based on the MM equation, they should be performed only when the MM equation is valid, that is, when the enzyme concentration is a much lower than the sum of the substrate concentration and the *K*_*M*_^7, 14^ (see below for more details). Because the value of *K*_*M*_ is usually not known a priori, to ensure the validity of the MM equation, *in vitro* experiments are typically performed with a much lower enzyme concentration than substrate concentration^15^. However, such conditions cannot be guaranteed *in vivo*, because endogenous enzyme concentrations are much higher than those used in a typical *in vitro* assay^16, 17^. It is therefore risky to use the MM equation to analyze *in vivo* data and to predict *in vivo* enzyme activity by using parameters estimated from an *in vitro* assay^15, 18^. Furthermore, even when the MM equation is valid, precise estimation is not guaranteed, because of the highly correlated structure and unidentifiability of the parameters^19–23^. That is, even though estimated parameters can fit the data accurately, the estimates can differ greatly from the actual values of *k*_*cat*_ and *K*_*M*_. Because of the identifiability issue, experimental designs to infer the maximum possible information about the parameters have been investigated^12, 13, 20–24^. For instance, to ensure that the parameters can be identified from the initial velocity assay, the initial concentration of substrate needs to be increased from a low level to a higher level until the reaction velocity is saturated. For the saturation, generally the initial substrate concentration needs to be larger than 10*K*_*M*_, but often such high concentrations cannot be achieved^24^. For the progress curve assay, the initial substrate concentration is recommended to be at a similar level to *K*_*M*_^23, 25^. Note that both assays require prior knowledge of *K*_*M*_, which gives rise to the conundrum that, in order to estimate *K*_*M*_, the approximate value of *K*_*M*_ needs to be known.

To overcome such limits on the inference using the model based on the MM equation, which is referred to as the sQ (standard QSSA) model, here we propose an alternative approach. In our approach, we use a different approximate model that is derived with the total QSSA and is referred to as the tQ (total QSSA) model^26–29^. By applying the Bayesian inference based on either the sQ or the tQ model to the product progress curve, we found that the estimates obtained with the sQ model were considerably biased when the enzyme concentration was not low. On the other hand, the estimates obtained with the tQ model were not biased for any combination of enzyme and substrate concentrations. Thus, with the tQ model, the experimental data from various conditions can be pooled without any restrictions to improve the accuracy and precision of the estimation. For instance, when two sets of timecourse data obtained under low and high enzyme concentrations are used together, the tQ model, but not the sQ model, leads to accurate and precise estimation. Another advantage of our approach is that, by analyzing the scatter plots of current estimates, the next optimal experiment to ensure the parameter identifiability can be easily designed without requiring any prior knowledge of the *k*_*cat*_ and *K*_*M*_ values. The proposed optimized design yields accurate and precise estimation from a minimal amount of data simulated based on the kinetics of various enzymes: chymotrypsin, fumarase and urease, which have disparate catalytic efficiencies (*k_cat_/K*_*M*_). We provide a publicly accessible computational package that performs the Bayesian inference based on the tQ model, thus leading to accurate and efficient estimation of enzyme kinetics.

## Results

### Two types of models describing enzyme kinetics: The sQ and tQ models

A fundamental enzyme reaction consists of a single enzyme and a single substrate, where the free enzyme (E) reversibly binds with the substrate (S) to form the complex (C), and the complex irreversibly dissociates into the product (P) and the free enzyme:

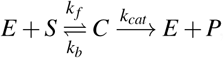

where the total enzyme concentration (*E_T_ ≡ C* + *E*) and the total substrate and product concentration (*S_T_ ≡ S* + *C* + *P*) are conserved. A popular model describing the accumulation of the product over time is based on the MM equation, as follows (see Supplementary Method for detailed derivation):

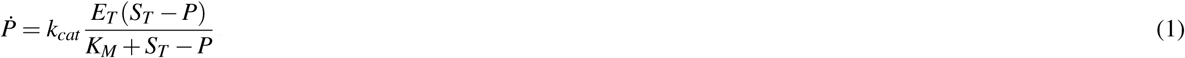

where *K*_*M*_ = (*k*_*b*_ + *k*_*cat*_)*/k*_*f*_ is the Michaelis-Menten constant and *k*_*cat*_ is the catalytic constant. This sQ model derived with the standard QSSA has been widely used to estimate the kinetic parameters, *K*_*M*_ and *k*_*cat*_ from the progress curve of the product^8–11, 23, 25^. Another model describing the accumulation of the product is derived with the total QSSA; it was developed later than the sQ model and thus has received less attention for parameter estimation^26–29^:

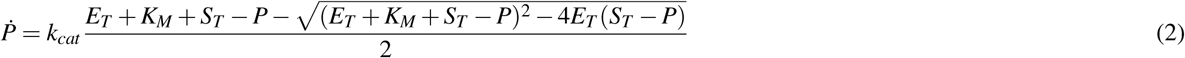

Although this tQ model is more complicated than the sQ model, it is accurate over wider ranges than the sQ model. Specifically, the sQ model is accurate when

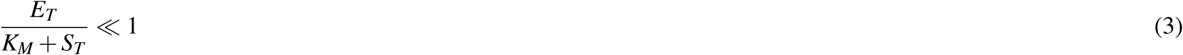

which requires a low enzyme concentration^7, 14^. On the other hand, the tQ model is accurate when

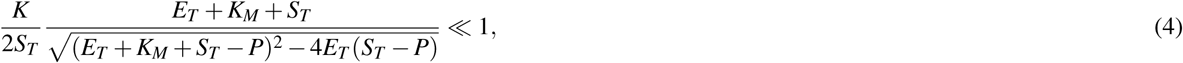

where *K* = *k_b_/k*_*f*_ is the dissociation constant^27–29^. Importantly, this condition is generally valid and thus the tQ model, unlike the sQ model, is accurate even when the enzyme is in excess. See^14, 30^ for more details.

We also investigated the accuracy of the stochastic simulations performed with both models. Specifically, we compared the stochastic simulations using the Gillespie algorithm based on the propensity functions from either the original full model (described in Table S1), the sQ model (Table S2), or the tQ model (Table S3) for 9 different conditions^31–36^: *E*_*T*_ is either lower than, similar to, or higher than *K*_*M*_, and *S*_*T*_ is also either lower than, similar to, or higher than *K*_*M*_ (Fig. 1). The stochastic simulations of the sQ model fail to approximate those of the original full model when *E*_*T*_ is not low (i.e., *E*_*T*_ is lower than neither *S*_*T*_ nor *K*_*M*_). On the other hand, stochastic simulations using the tQ model are accurate for all conditions (Fig. 1), as is consistent with a recent study showing that stochastic simulations with the sQ and the tQ models are accurate when their deterministic validity conditions hold (Eqs. (3) and (4))^37, 38^. Taken together, the tQ model is valid for a wider range of conditions than the sQ model is in both the deterministic and the stochastic sense.

**Figure 1.**
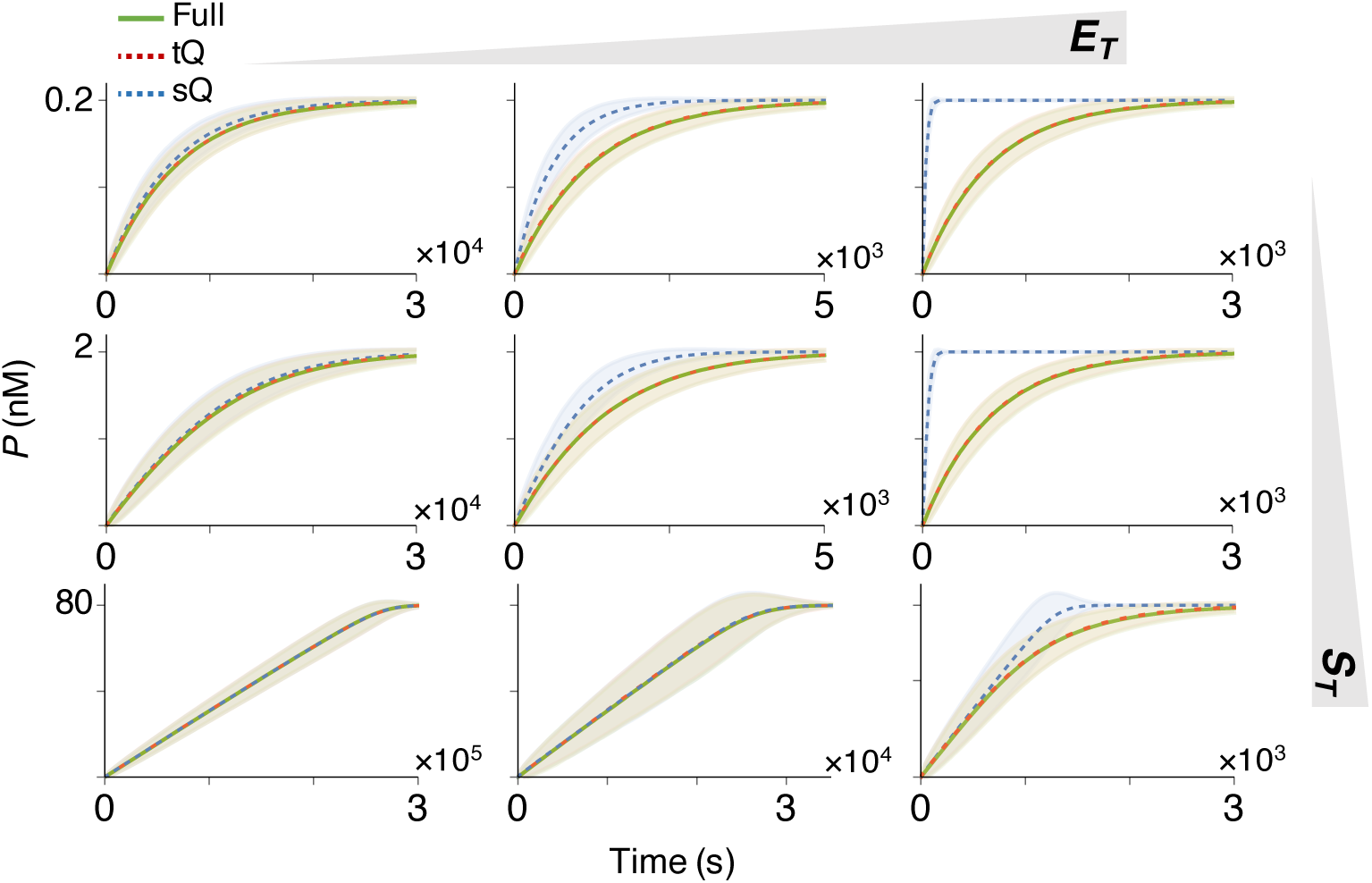
Whereas the sQ model fails to approximate the original full model as *E*_*T*_ increases, the tQ model is accurate regardless of *E*_*T*_. Stochastic simulations of the original full model (Table S1), the sQ model (Table S2), and the tQ model (Table S3) were performed with *S*_*T*_ = 0.2, 2, or 80nM, and *E*_*T*_ = 0.2, 2, or 40nM. Note that these concentrations are either lower than, similar to, or higher than *K_M_ ≈*2nM. Here, the lines and colored ranges represent a mean trajectory and fluctuation range (*±*2*σ* from the mean) of 10^4^ stochastic simulations.

### Estimation with the tQ model is unbiased for any combination of enzyme and substrate concentrations

Because the tQ model is accurate for a wider range of conditions than the sQ model is (Fig. 1), we hypothesized that the parameter estimation based on the tQ model is also accurate for more general conditions. To investigate this hypothesis, we first generated 10^2^ noisy progress curves of *P* from the stochastic simulations of the original full model (Fig. S1). Then, we inferred parameters (*k*_*cat*_ and *K*_*M*_) from these simulated data sets by applying the Bayesian inference with the likelihood functions based on either the sQ or the tQ model, under weakly informative gamma priors (Fig. S2) (see Methods for details). Note that throughout this study, we have used the simulated product progress curves (e.g. Fig. S1) because we need to know the true values of parameters for the accurate comparison of the estimations based on the sQ model and the tQ model.

We first focused on the estimation of the *k*_*cat*_ under the assumption that the value of *K*_*M*_ is known. When *E*_*T*_ is low, so that both the sQ and the tQ models are accurate (Fig. 1 left), posterior samples obtained with both models are similar and successfully capture the true value of *k*_*cat*_ (Fig. 2a left). The posterior samples obtained with the two models are similar because, when *E*_*T*_ is low and thus *E_T_ ≪ S*_*T*_ + *K*_*M*_, both models (Eqs. 1 and 2) are approximately equivalent as follows:

**Figure 2.**
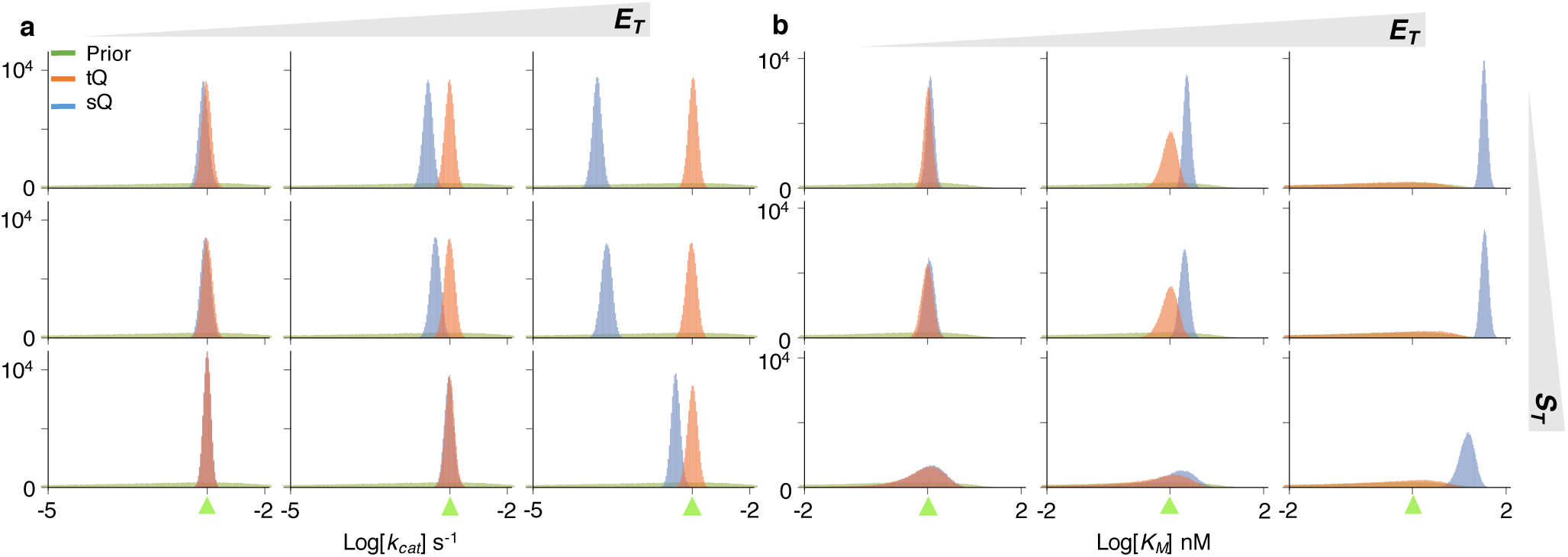
The estimation of a single parameter (*k*_*cat*_ or *K*_*M*_) with either the sQ or the tQ model. For each condition (*S*_*T*_ = 0.2, 2, or 80nM, and *E*_*T*_ = 0.2, 2, or 40nM), 10^5^ posterior samples of either *k*_*cat*_ (a) or *K*_*M*_ (b) were obtained by applying the Bayesian inference to 10^2^ noisy data sets (Fig. S1) (see Methods for details). When the *k*_*cat*_ is sampled, the *K*_*M*_ is fixed at its true value (a) and vice versa (b). Here, green triangles indicate the true values of the parameters. Whereas the estimates of *k*_*cat*_ and *K*_*M*_ obtained with the sQ model are biased as *E*_*T*_ increases, those obtained with the tQ model have negligible bias regardless of conditions (See Fig. S3 for box plots of estimates). As *E*_*T*_ or *S*_*T*_ increases, the posterior variance of *K*_*M*_ increases when the tQ model is used.

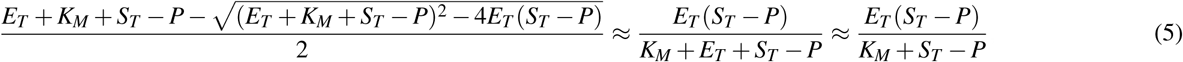

where the first approximation comes from the Taylor expansion in terms of *E*_*T*_ (*S_T_ − P*)*/*(*E*_*T*_ + *K*_*M*_ + *S_T_ P*) ≪ 1 (see^27–29^ for details). Therefore, when *E_T_ S*_*T*_ + *K*_*M*_ and thus the sQ model is accurate, estimations with the sQ and the tQ models should be similar. On the other hand, when *E*_*T*_ is high, they show clear differences (Fig. 2a right): the posterior samples obtained with the sQ model show large errors, while those obtained with the tQ model accurately capture the true value of *k*_*cat*_.

Similar results are also observed in the box plots of posterior means and posterior coefficient of variations (CVs) (Fig. S3a and b). Whereas posterior means obtained with the sQ model are biased when *E*_*T*_ is high, those obtained with the tQ model are accurate for all conditions (Fig. S3a). In particular, narrow distributions of posterior means indicate that the estimation of *k*_*cat*_ with the tQ model is robust aginst the noise in the data (Fig. S1). Furthermore, posterior CVs are much smaller than prior CVs (Fig. S3b), indicating precise estimation of *k*_*cat*_ with the tQ model.

Next, *K*_*M*_ was estimated under the assumption that the value of *k*_*cat*_ is known (Fig. 2b). Posterior samples of the *K*_*M*_ obtained with the sQ model again show errors that grow with increasing *E*_*T*_. Note that the estimates of the *K*_*M*_ are biased upward, which implies that using the posterior estimates of *K*_*M*_ to validate the MM equation (*K_M_ ≫ E_T_*) can be misleading. On the other hand, the estimates of *K*_*M*_ obtained with the tQ model are little biased for all conditions. However, unlike the narrow posterior distributions of *k*_*cat*_ (Fig. 2a), those of *K*_*M*_ obtained with the tQ model become wider; so precision decreases as *E*_*T*_ or *S*_*T*_ increases (Fig. 2b). These patterns are also observed in the box plots of posterior means and posterior CVs (Fig. S3c and d). The identifiability problem arises because, when *E_T_ ≫ K*_*M*_ or *S_T_ ≫ K*_*M*_ and thus *E*_*T*_ + *S_T_ ≫ K*_*M*_, the *K*_*M*_ is negligible in the tQ model (Eq. 2), as follows:

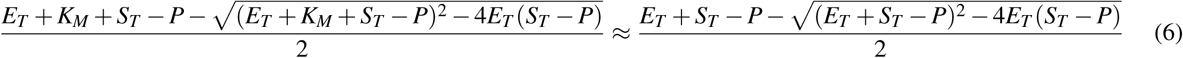

Specifically, when *K*_*M*_ is too low, the value of *K*_*M*_ has little effect on the dynamics of the tQ model and thus the *K*_*M*_ is structurally unidentifiable. Taken together, the estimations of *K*_*M*_ with both the sQ and the tQ models are not satisfactory, although for different reasons: estimations with the sQ model can be biased and those with the tQ model can be structurally unidentifiable (Fig. 2b). Similar patterns were also observed when a more informative prior was given (Fig. S4). In particular, even with the informative prior, estimates obtained with the sQ model still show considerable error as *E*_*T*_ increases.

### Simultaneous estimation of *k*_*cat*_ **and** *K*_*M*_ suffers from the lack of identifiability

Next, we considered simultaneous estimation of two parameters, *k*_*cat*_ and *K*_*M*_, which is the typical goal of enzyme kinetics. For the same gamma priors used in the single-parameter estimation (Fig. 2), the distributions of posterior samples obtained with both models became wider overall (Fig. 3). To find the reason for such imprecise estimation, we analysed the scatter plots of posterior *k*_*cat*_ and *K*_*M*_ samples (Fig. 4). When *S_T_ ≪ K*_*M*_ (Fig. 4 a-c), the posterior samples of *k*_*cat*_ and *K*_*M*_ obtained with the sQ model exhibited a strong correlation, because the dynamics of the sQ model depend only on the ratio *k*_*cat*_ /*K*_*M*_, as seen in the following approximation:

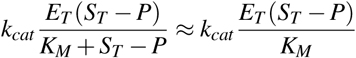

**Figure 3.**
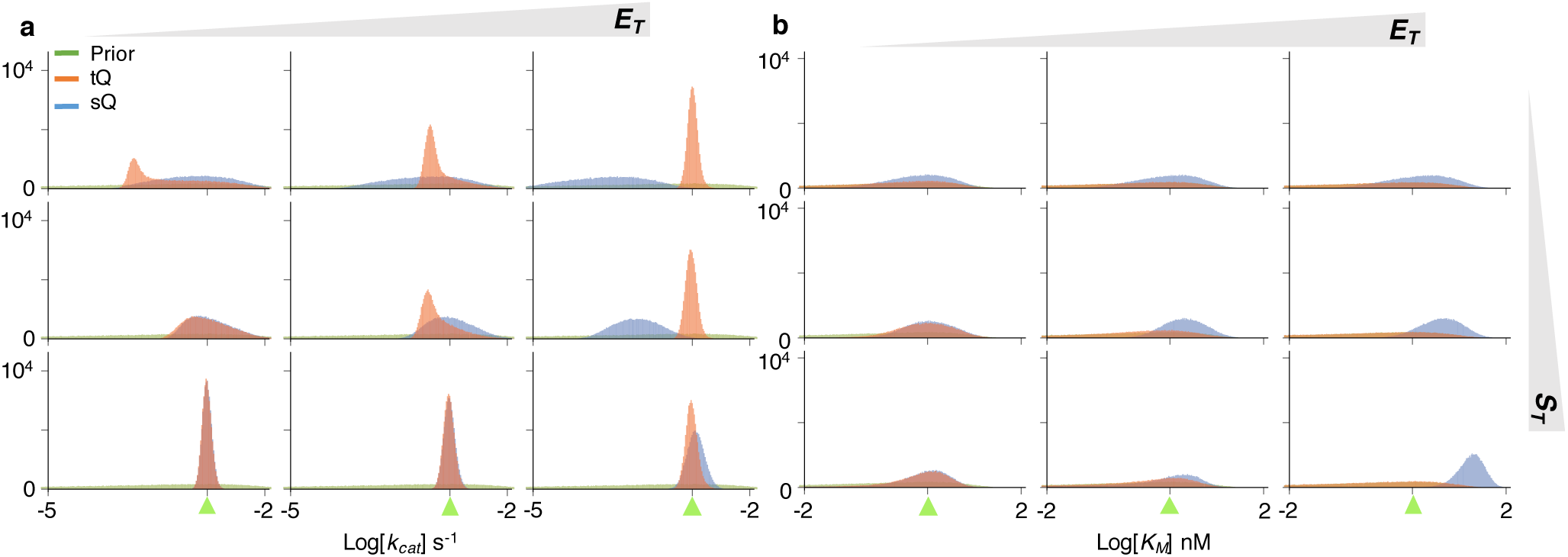
Simultaneous estimation of two parameters (*k*_*cat*_ and *K*_*M*_) with either the sQ or the tQ model. From the same 10^2^ data sets (Fig. S1) used in the single-parameter estimation (Fig. 2), 10^5^ posterior samples of the *k*_*cat*_ (a) and the *K*_*M*_ (b) were obtained together. Although the same prior is given, the posterior distributions become wider than the single-parameter estimation (Fig. 2). Here, green triangles indicate the true values of *k*_*cat*_ or *K*_*M*_.

**Figure 4.**
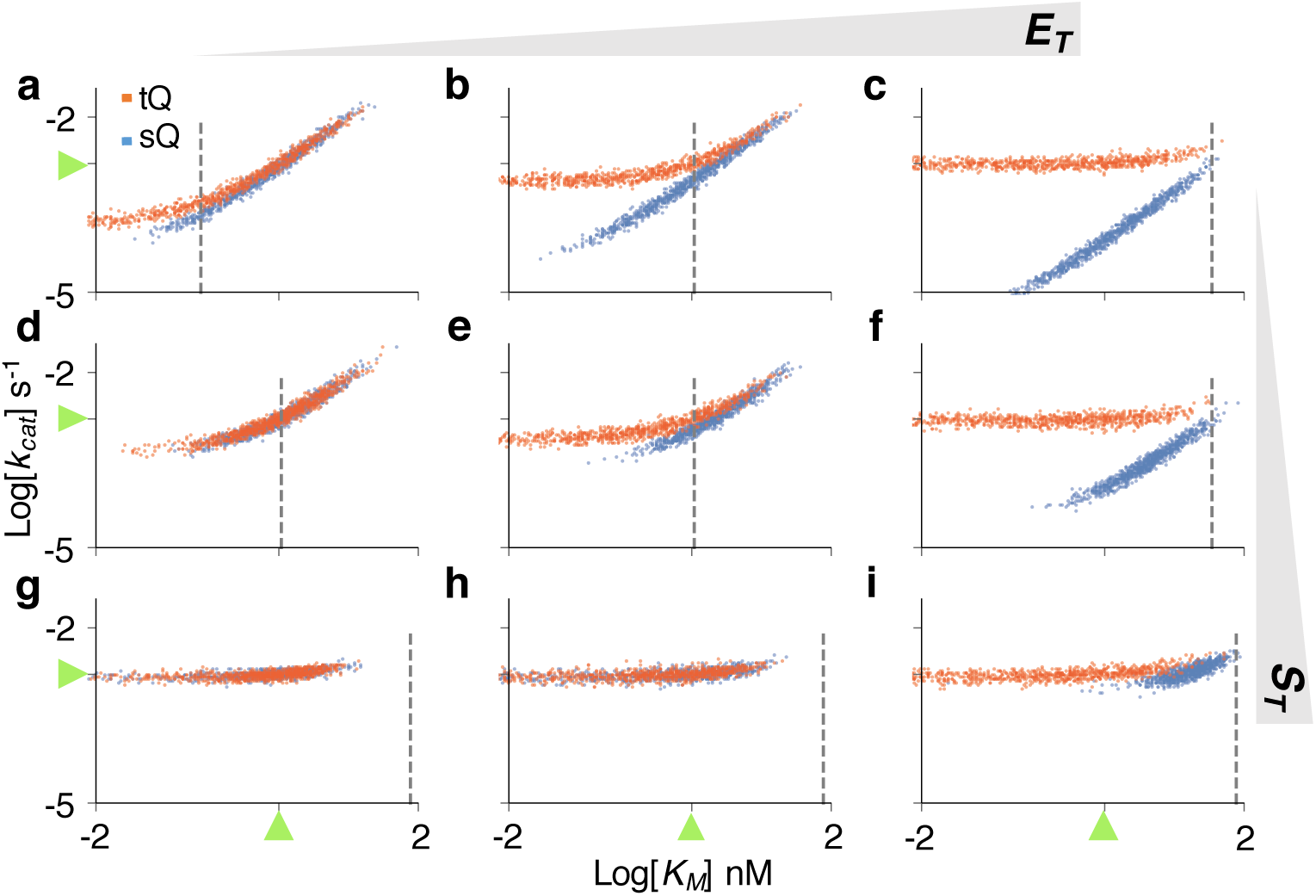
The scatter plots of posterior samples obtained with the two-parameter estimation (Fig. 3). The scatter plots imply two types of structure unidentifiability: strong correlation between *k*_*cat*_ and *K*_*M*_, and unidentifiability of *K*_*M*_, which is represented as a horizontal plot. Positively correlated scatter plots of the tQ model are changed to horizontal ones when the sampled *K*_*M*_ is much lower than *S*_*T*_ + *E*_*T*_ (dashed gray lines). Here, green triangles represent the true values of parameters.

where *K*_*M*_ ≫ *S*_*T*_ ≥ *S*_*T*_ − *P* is used. On the other hand, when *S*_*T*_ ≫ *K*_*M*_ (Fig. 4g-i), the scatter plot of the sQ model becomes horizontal, indicating the structure unidentifiability of the *K*_*M*_. Indeed, the value of *K*_*M*_ has nearly no effect on the dynamics of the sQ model, as seen in the following approximation:

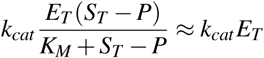

where *K*_*M*_ + *S*_*T*_ ≈ *S*_*T*_ is used as *S*_*T*_ ≫ *K*_*M*_. Such lack of parameter identifiability when *S*_*T*_ ≪ *K*_*M*_ or *S*_*T*_ ≫ *K*_*M*_ is consistent with previous studies, which recommend using *S_T_ ≈ K*_*M*_ for more precise estimation^22, 23^. However, even when *S_T_ ≈ K*_*M*_, estimates are still imprecise (Fig. 3 middle). Furthermore, as *E*_*T*_ increases, the sQ model is biased like the single-parameter estimation. Based on this analysis it appears that the simultaneous estimation of *k*_*cat*_ and *K*_*M*_ with the sQ model is challenging because of both identifiability and bias problems.

When *E*_*T*_ ≫ *K*_*M*_ or *S*_*T*_ ≫ *K*_*M*_, the *K*_*M*_ has a negligible effect on the dynamics of the tQ model (Eq. 6), and thus only *k*_*cat*_ was identifiable in the single-parameter estimation (Fig. 2 right or bottom). Similarly, when both *k*_*cat*_ and *K*_*M*_ are inferred simultaneously with the tQ model, estimation of only *k*_*cat*_ is accurate and precise (Fig. 3 right or bottom), as is shown by the horizontal scatter plots along the true value of *k*_*cat*_ (Fig. 4 c, f, g-i). In other cases (when neither *E*_*T*_ ≫ *K*_*M*_ nor *S_T_ ≫ K_M_*), posterior variance of both parameters dramatically increases compared to the single-parameter estimation (Fig. 2 and 3 left and top). Such imprecise estimation stems from two sources, according to the scatter plots (Fig. 4a,b,d,e). When *k*_*cat*_ and *K*_*M*_ decrease together, the behavior of the tQ model changes little, which leads to the strong correlation between posterior samples of *k*_*cat*_ and *K*_*M*_. As the estimates of *K*_*M*_ keep decreasing together with those of *k*_*cat*_, so that they become much less than *E*_*T*_ + *S*_*T*_ (dashed vertical line of Fig. 4), the tQ model no longer depends on the value of *K*_*M*_, as shown in Eq. 6, and thus the scatter plots become horizontal.

### Combined data from different experiments allow accurate and precise estimation with the tQ model

As shown above, the estimation of both *k*_*cat*_ and *K*_*M*_ using a single progress curve suffers from considerable bias and lack of identifiability (Figs. 3 and 4), as is consistent with previous studies reporting that a progress curve obtained from a single experiment is not enough to identify both parameters simultaneously^19^. Thus, here, we investigate whether using multiple timecourse data sets obtained under different experimental conditions can improve the estimation.

In typical *in vitro* assays, progress curves are measured with either a fixed *S*_*T*_ and varied *E*_*T*_ or a fixed *E*_*T*_ and varied *S*_*T*_ ^8–11, 39^. We first consider the case when progress curves are measured with a fixed *S*_*T*_ and a varied *E*_*T*_. Specifically, progress curves from both low and high *E*_*T*_ are used to estimate parameters for a fixed *S*_*T*_ at different levels (Fig. S1 top and bottom). In this case, posterior samples obtained with the sQ model show considerable errors as the data from high *E*_*T*_ is used (Figs. 5a and S5). On the other hand, the posterior samples obtained with the tQ model accurately capture the true values of both *k*_*cat*_ and *K*_*M*_ with low variance (Figs. 5a and S5). Such improvement stems from the fact that data obtained under the low and high *E*_*T*_ provide different types of information for parameter estimation. Specifically, from the high *E*_*T*_ data, although the *K*_*M*_ is not identifiable, the *k*_*cat*_ can be accurately estimated with the tQ model (Fig. 4c, f, i). Such accurate estimation of *k*_*cat*_ from the high *E*_*T*_ data can prevent the correlation between the *k*_*cat*_ and the *K*_*M*_ when they are estimated from the low *E*_*T*_ data (Fig. 4a, d). Indeed, the narrow scatter plots of the tQ model (Fig. 5b) are the intersection of two scatter plots, a horizontal one obtained with the high *E*_*T*_ data (Fig. 4 c, f) and a nonhorizontal one obtained with the low *E*_*T*_ data (Fig. 4 a, d). However, when *S*_*T*_ is high, the scatter plot from the low *E*_*T*_ also becomes horizontal (Fig. 4c), and thus the synergistic effect of using combined data decreases (Fig. 5a and b right). Taken together, the tQ model can accurately estimate both parameters from the combination of low *E*_*T*_ and high *E*_*T*_ data when *S*_*T*_ is not much larger than *K*_*M*_. Note that such low *S*_*T*_ is preferred for *in vitro* experiments^24, 39–41^ and is the case for most physiological conditions^24^.

**Figure 5.**
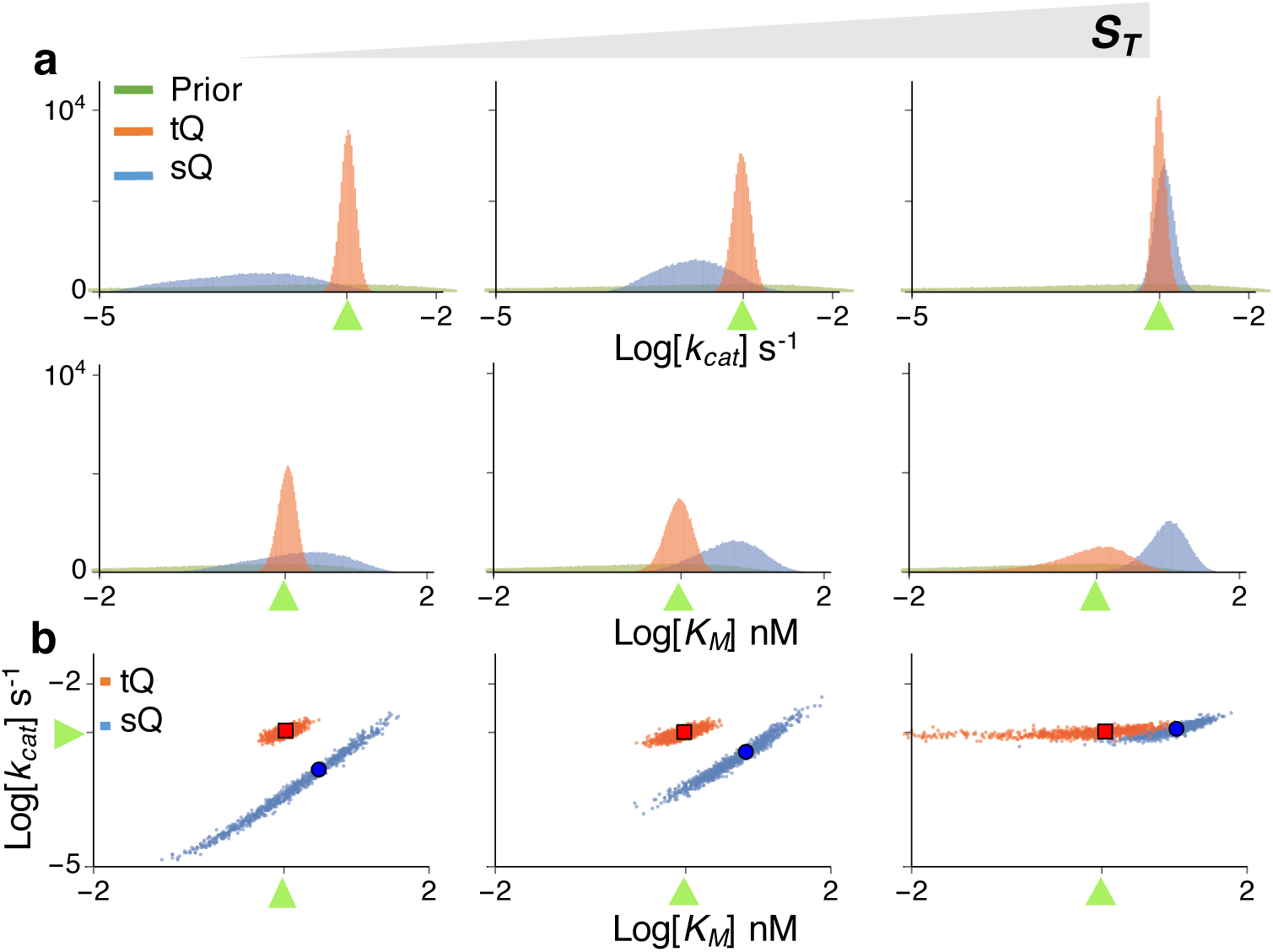
When data obtained under low *E*_*T*_ and high *E*_*T*_ are used together, the accuracy and precision of estimaties obtained with the tQ model, but not with the sQ model, are enhanced. (a) Posterior samples are inferred using data sets from *E*_*T*_ = 0.2nM (Fig. S1 top) and *E*_*T*_ = 40nM (Fig. S1 bottom) together for either *S*_*T*_ = 0.2, 2, or 80nM. The posterior variance of the tQ model dramatically decreases to the level of the single-parameter estimation (Fig. 2). However, the estimates of the sQ model show considerable bias. Here, green triangles represent the true values of *k*_*cat*_ or *K*_*M*_. (b) The scatter plots of the posterior samples. Here green triangles, blue circles, and red squares represent true values, posterior means of the sQ model, and those of the tQ model, respectively.

Next, we consider the case when progress curves are measured with a fixed *E*_*T*_ and a varied *S*_*T*_. Specifically, the combination of two progress curves from low and high *S*_*T*_ is used to infer parameters for a fixed *E*_*T*_ at different levels (Fig. S1 left and right). When *E*_*T*_ is low, and thus the sQ and the tQ models behave similarly (Eq. 5), posterior samples obtained with both models accurately capture the true values of *k*_*cat*_ and *K*_*M*_ (Figs. 6a left and S6). Again, the narrow scatter plot (Fig. 6b left) is obtained as the intersection of a nonhorizontal scatter plot of low *S*_*T*_ (Fig. 4a) and a horizontal scatter plot of high *S*_*T*_ (Fig. 4g). However, as *E*_*T*_ increases, and thus the sQ model becomes less accurate, those obtained with the sQ model are biased, as expected (Figs. 6a right and S6). Whereas such biases are not observed in those obtained with the tQ model, the precision of *K*_*M*_ estimates decreases as *E*_*T*_ increases, as in the single-parameter estimation (Fig. 2 and Eq. 6).

**Figure 6.**
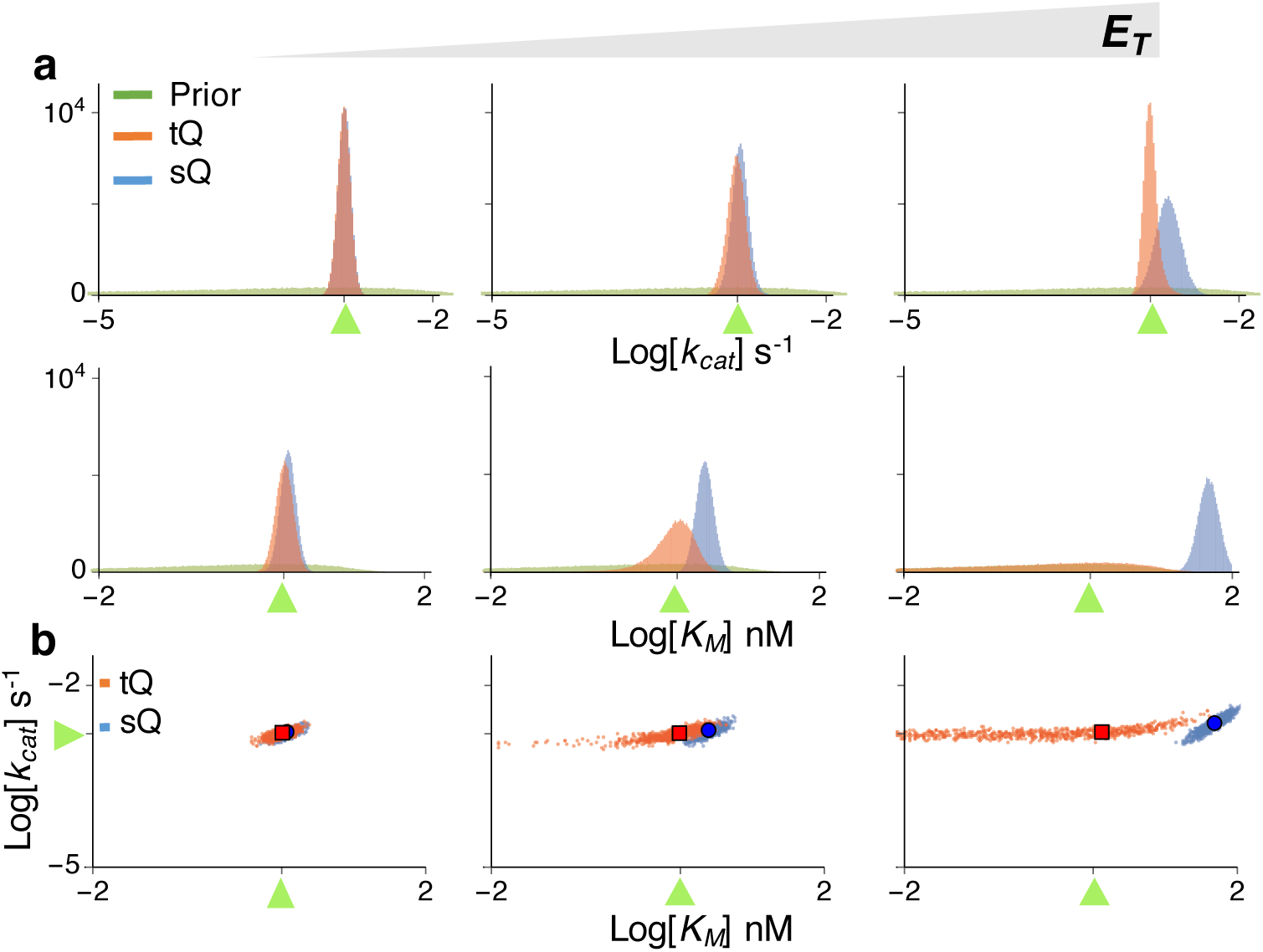
Estimation using the data obtained under low *S*_*T*_ and high *S*_*T*_ together. (a) Posterior samples are inferred using data sets from *S*_*T*_ = 0.2nM (Fig. S1 left) and *S*_*T*_ = 80nM (Fig. S1 right) together for either *E*_*T*_ = 0.2, 2, or 40nM. When *E*_*T*_ is low, both the sQ and the tQ models allow accurate and precise estimation. As *E*_*T*_ increases, the estimates obtained with the sQ model become inaccurate, and the estimates of *K*_*M*_ obtained with the tQ model become less precise, similar to the single-parameter estimation (Fig. 2). Here, green triangles represent the true values of *k*_*cat*_ or *K*_*M*_. (b) The scatter plots of the posterior samples. Here green triangles, blue circles, and red squares represent true values, posterior means of the sQ model, and those of the tQ model, respectively.

### Optimal design of experiments for accurate and efficient estimation with the tQ model

When a progress curve obtained from a single experiment is used, the posterior scatter plots of the tQ model can be categorized as a correlated type (Fig. 4a,b,d,e) and a horizontal type (Fig. 4c, f, g-i). The intersections of these two different types of scatter plots tend to be narrowly distributed near the true value (Figs. 5b and 6b). Thus, combining two such data sets allows accurate estimation of both *k*_*cat*_ and *K*_*M*_ (Figs. 5a and 6a). Specifically, a progress curve measured under *E*_*T*_ ≪ *K*_*M*_ and *S*_*T*_ ≪ *K*_*M*_ (Fig. 4a,b,d,e) and one measured under *E*_*T*_ ≫ *K*_*M*_ or *S*_*T*_ ≫ *K*_*M*_ (Fig. 4c, f, g-i) provide different types of information for parameter estimation; so using both data sets leads to successful estimation. However, it is hard to compare the values of *S*_*T*_, *E*_*T*_, and *K*_*M*_ in practice, because the value of *K*_*M*_ is usually unknown a priori. This problem can be easily resolved by using the scatter plot. That is, if the posterior scatter plot obtained from the first experiment is horizontal, then both *E*_*T*_ and *S*_*T*_ should be decreased for the next experiment, so that the nonhorizontal scatter plot can be obtained (Fig. 7a). On the other hand, if the scatter plot from the first experiment shows a strong correlation between *K*_*M*_ and *k*_*cat*_, then either *S*_*T*_ or *E*_*T*_ should be increased in the next experiment (Fig. 7b). Basically, without any prior information of the value of *K*_*M*_ and *k*_*cat*_, the shape of the scatter plots of the current estimates determines the next optimal experimental design, which ensures accurate and precise estimation. However, this approach cannot be used with the sQ model, because estimation with the sQ model can be biased, depending on the relationship between *E*_*T*_ or *S*_*T*_ and *K*_*M*_, which is unknown a priori. That is, unlike the tQ model, precise estimation does not always guarantee accurate estimation with the sQ model, as seen above (e.g. Fig 5a right).

**Figure 7.**
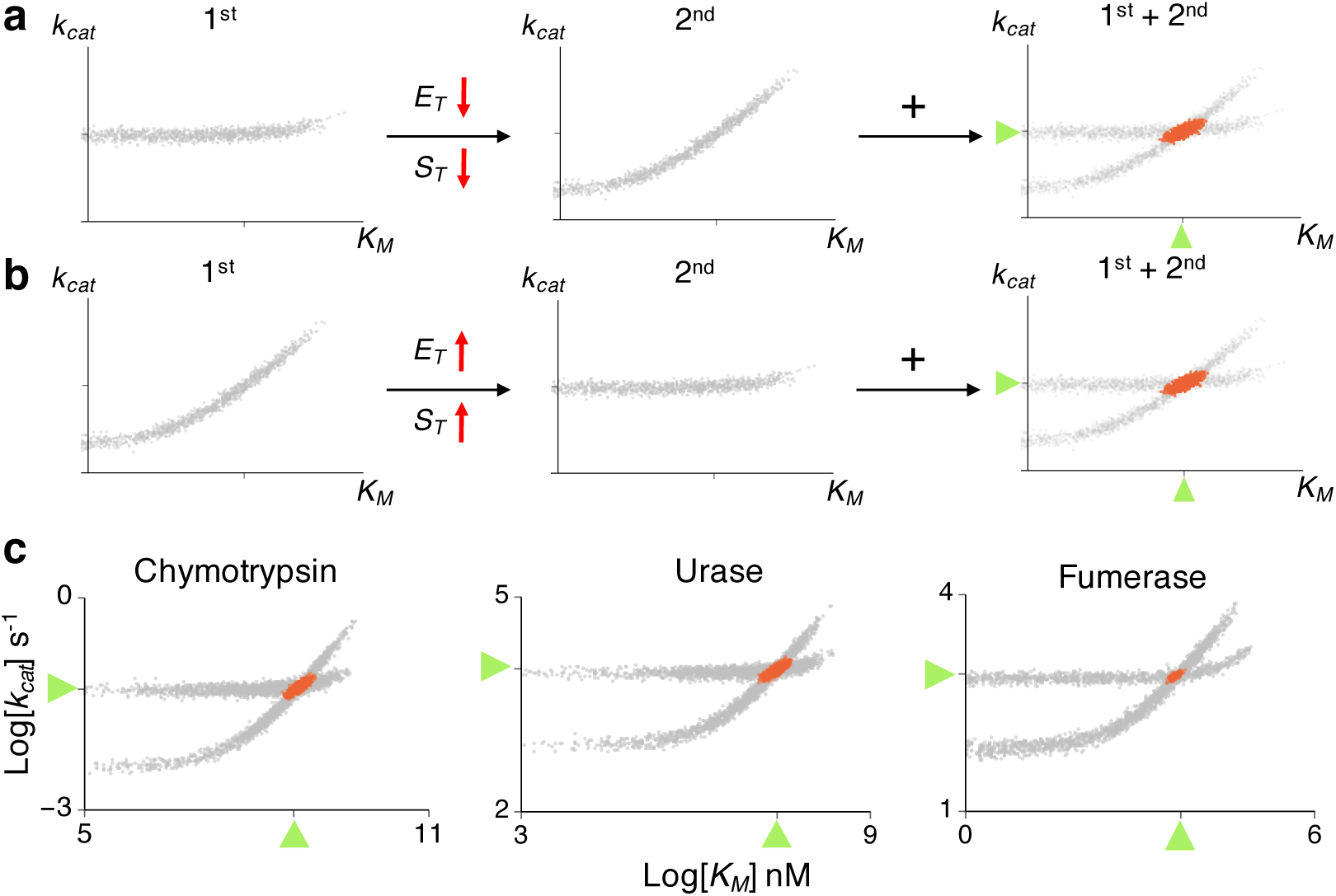
The optimal experimental design for accurate and precise estimation with the tQ model. (a) When the scatter plot of posterior samples from the first experiment is horizontal, *E*_*T*_ and *S*_*T*_ need to be decreased to obtain the nonhorizontal scatter plot in the next experiment. Then, using the combination of the two experiments leads to accurate and precise estimation (red scatter plots). (b) When the scatter plot from the first experiment is nonhorizontal, *E*_*T*_ or *S*_*T*_ need to be increased in the next experiment to obtain a horizontal scatter plot. (c) Inference with a single progress curve from the low *E*_*T*_ (0.1*K*_*M*_) and the high *E*_*T*_ (10*K*_*M*_) leads to nonhorizontal and horizontal scatter plots, respectively, for chymotrypsin, urease, and fumarase (gray scatter plots). When both data sets were used together, accurate estimates were obtained for all enzymes (red scatter plots). Here, low *S*_*T*_ (0.1*K*_*M*_) is used. Here, green triangles represent the true values of the parameters.

We test whether the proposed approach with the tQ model can accurately estimate *k*_*cat*_ and *K*_*M*_ for catalysis of the N-acetylglycine ethyl ester, fumarate, and urea by the enzymes the chymotrypsin, urease, and fumarase, respectively (Fig. 7c). These three enzymes were chosen because they have disparate catalytic efficiencies (*k_cat_ /K_M_*)^1^: 0.12, 4 10^5^, and 1.6 10^8^*s*^*−1*^*M*^*−1*^, respectively. For each enzyme, 10^2^ noisy timecourse data sets were generated using stochastic simulations based on known enzyme kinetic parameters^1^. When progress curves obtained with low *E*_*T*_ and low *S*_*T*_ are used, as expected, nonhorizontal scatter plots of posterior samples were obtained for all three enzymes (Fig. 7c). This indicates that either *E*_*T*_ or *S*_*T*_ should be increased in the next experiment to obtain a horizontal scatter plot. Indeed, when a progress curve with a 100-fold increase of *E*_*T*_ was used, horizontal scatter plots were obtained for all enzymes (Fig. 7c). Therefore, when these two progress curves are used together, both *k*_*cat*_ and *K*_*M*_ can be accurately estimated (Fig. 7c red dots). These results support that such two-step optimized experimental design (Fig. 7a and b) to get two different types of scatter plots allows accurate and efficient estimation of enzyme kinetics with the tQ model.

## Discussion

The standard approach for estimating enzyme kinetic parameters even today continues to be based on the 100-year old MM equation (Eq. 1)^5, 6^. However, when enzyme concentration is high, this approach can lead to biased estimation (Fig. 2). Even when enzyme concentration is relatively low, it may not be possible to identify kinetic parameters (Fig. 3 and 4). To overcome the limitations of the canonical approach, we proposed an estimation method based on an alternative to the MM equation: the tQ model (Eq. 2), which is derived with the total QSSA^26–29^. Because the estimation procedure with the tQ model is not biased regardless of enzyme or substrate concentrations (Fig. 2), more accurate and precise estimations can be made when pooled data from different experimental conditions are used, unlike the canonical approach (Figs. 5 and 6). It appears thus that the tQ model is especially appropriate for creating a consistent Bayesian inferential framework, which becomes more accurate as more data is used.

The canonical enzyme kinetic assay based on the MM equation generally requires a large excess of substrate over enzyme^42^. However, such conditions impose experimental limitations and cannot be always guaranteed and verified^15^. For instance, it is hard to generate a high concentration of barely soluble substrate^24^, and a low concentration of substrate is required for sensitive kinetic analysis, e.g., in the case of QD-FRET-based probes^39–41^. Importantly, to analyze *in vivo* enzyme kinetics, where enzyme concentration is often high^16–18^, our approach, but not the canonical approach, can be used. For example, one needs to estimate the kinetic parameters underlying drug metabolism by CYP enzymes in the liver in order to predict the effects of drugs, as is essential for drug development^43^. Because of dosing requirements for potent drugs, the amount of CYP enzyme can greatly exceed the drug amount in the liver^44, 45^. Another large area where our estimation method can be applied is in the development of nanobiosensors, which measure *in vivo* activity of a specific enzyme for precise diagnostics, because such enzymes are often in large excess over biosensors^46, 47^.

KinTek Explorer has been widely used to estimate enzyme kinetic parameters from the progress curves^48–50^. This software provides the confidence contours, which reveal the relationships between the estimated parameters. This approach recommends using multiple data sets to narrow down the confidence contours and thus improve precision of estimates and resolve the unidentifiability issue. Our finding (Fig. 7a and b) can provide the specific type of data sets required for the identifiability of *k*_*cat*_ and *K*_*M*_, so that the KinTek Explore could perform parameter estimation more efficiently for the Michales-Menten type of enzyme reactions.

Since the initial velocity estimation with the MM equation is not accurate when enzyme concentration is high (Fig. 1), the standard initial velocity based on the MM equation would also be inaccurate^15^. On the other hand, the tQ model accurately captures the initial velocity for all conditions, and thus the modified initial velocity assay based on the tQ model is likely to be accurate over a wider range of conditions. To simplify such estimation procedures, an interesting future study could derive an analogous Lineweaver-Burk plot or the Hanes-Woolf plot^8, 9, 15, 42^ for the tQ model.

Even with relatively large noise in the data (Fig. S1), our proposed method leads to accurate estimation (Figs. 5-7), indicating its robustness against experimental noise and some minor inaccuracy of the tQ model in certain ranges of parameter observed in^14, 30^. Furthermore, if there are departures from simple non-inhibitory enzyme kinetics (e.g. inhibition of enzyme by product)^51, 52^, our method can be easily adjusted by modifying the tQ model (see^53, 54^ for the tQ model for other enzyme kinetics). Our work can also be used to improve the estimation of the kinetics underlying diverse biological functions, such as gene regulation^55, 56^, cellular rhythms^57–59^, quorum sensing^60, 61^, signal cascade^62, 63^ and membrane transport^64, 65^, where the MM equation has been widely used.

## Methods

### Simulated Data

To obtain timecourse data (Fig. S1) for Bayesian inference, stochastic simulations of the original full model (Table S1) were performed with the Gillespie algorithm^66^. *E*(0) = *E*_*T*_, *S*(0) = *S*_*T*_, *C*(0) = 0, and *P*(0) = 0 are used as initial conditions following the typical *in vitro* enzyme kinetics protocol.

### Description of the Bayesian inference approach

The Bayesian inference approach is used to estimate the catalytic constant *k*_*cat*_ and the Michaelis-Menten constant *K*_*M*_, based on the hazard function, with respective rates described in Eq. 1 for the sQ model and in Eq. 2 for the tQ model (Fig. S2). The likelihood functions are constructed based on an approximation to the underlying Markov model^66^ as follows.

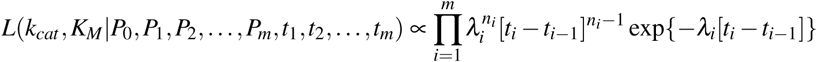

where *λ*_*i*_ is given by

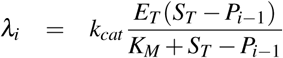, for the sQ model
or

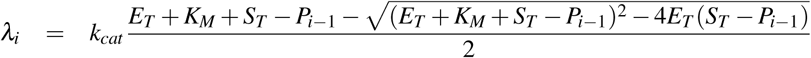, for the tQ model,
where *P*_*i*_ is the scaled number of product molecules observed at time point *t*_*i*_ over [0, T] = [*t*_*0*_, *t*_*m*_] and *n*_*i*_ = *P*_*i*_ − *P*_*i*_ − _1_ is an observed increment of *P*_*i*_. With these likelihood functions, the usual independent gamma priors^67^ are assigned to *k*_*cat*_ and *K*_*M*_ in order to get their posterior distributions with the help of the Markov Chain Monte Carlo (MCMC) method. Weakly informative gamma priors are used for both *k*_*cat*_ and *K*_*M*_: their prior means are the same as their true values, and their prior variance is 10 times larger than the prior mean, which covers orders of magnitude (e.g. Fig. 2). The estimation of a single parameter, i.e., either *k*_*cat*_ or *K*_*M*_ is done conditionally on the other parameter. For estimating the two parameters simultaneously, the Gibbs sampler method is used. In order to draw the sample for *K*_*M*_, we also use the Metropolis-Hastings algorithm within the Gibbs sampler step. See Supplementary material for further details.

### Computational code

The R package that performs the Bayesian inference based on the tQ model is available on the CRAN repository (www&.).

## Acknowledgements

The research was initiated when all authors were visiting The Mathematical Biosciences Institute at the Ohio State University. This work was supported by Korea University Grant (BC), US National Science Foundation (NSF) Grant DMS-1440386 (GAR) and DMS-1318886 (GAR), the National Research Foundation of Korea grant N01160447 (JKK), and the TJ Park Science Fellowship of POSCO TJ Park Foundation (JKK).

## Author contributions statement

JKK designed the research. BC and JKK performed simulations and analysis. All authors discussed the results and wrote the manuscript.

## Competing interests

The authors declare that they have no competing interests.

